# Exploring Applications of Crowdsourcing to Cryo-EM

**DOI:** 10.1101/220145

**Authors:** Jacob Bruggemann, Gabriel C. Lander, Andrew I. Su

## Abstract

Extraction of particles from cryo-electron microscopy (cryo-EM) micrographs is a crucial step in processing single-particle datasets. Although algorithms have been developed for automatic particle picking, these algorithms generally rely on two-dimensional templates for particle identification, which may exhibit biases that can propagate artifacts through the reconstruction pipeline. Manual picking is viewed as a gold-standard solution for particle selection, but it is too time-consuming to perform on data sets of thousands of images. In recent years, crowdsourcing has proven effective at leveraging the open web to manually curate datasets. In particular, citizen science projects such as Galaxy Zoo have shown the power of appealing to users' scientific interests to process enormous amounts of data. To this end, we explored the possible applications of crowdsourcing in cryo-EM particle picking, presenting a variety of novel experiments including the production of a fully annotated particle set from untrained citizen scientists. We show the possibilities and limitations of crowdsourcing particle selection tasks, and explore further options for crowdsourcing cryo-EM data processing.

## 1. Introduction

In the past several years cryo-electron microscopy (cryo-EM) has become a powerful tool for elucidating the structures of macromolecular complexes to near-atomic resolution, and has been effectively used to solve structures of membrane-bound and non-rigid proteins that are difficult to crystallize. Handling low signal-to-noise ratio cryo-EM data necessitates processing large amounts of data, involving thousands of individual micrographs each containing hundreds of particles. A crucial, early step in cryo-EM processing is the selection of individual protein particles from EM micrographs to be used in generating a 3D reconstruction. In the past, particles were hand-picked by a researcher after data collection, but since cryo-EM datasets can now consist of thousands of micrographs and hundreds of thousands of particles, manual picking has become viewed as an unnecessarily banal and time-consuming task for cryo-EM researchers [1].

As a result, many algorithms have been developed to automate particle picking and reduce the time required for this crucial step in EM processing. Popular methods either identify features common to particles, such as particle size with DoG Picker, or use supplied templates to identify similar-looking subsections of a micrograph [2]. Automatic methods are limited, however, in their ability to distinguish noise and contaminants from legitimate particles, and will sometimes misplace the center of particles in cases where they are closely packed. Inaccuracies in the collection of particle data can disrupt processing; in the challenging reconstruction of the HIV-1 envelope glycoprotein complex by Liao et al., Henderson noted that a lack of validation of the particle set picked via a template method begat a set of particles containing significant white noise, which nonetheless sufficiently matched the templates provided [3, 4]. Indeed, manual selection by a trained microscopist is still viewed as an ideal strategy in many cases, especially when templates are not available or the protein particles are ill-defined in the micrographs. Implementing manual selection necessitates an immense amount of time and effort for this single processing step; as an example, Fan et al. manually boxed out 156,805 particles from 3,743 micrographs when determining the structure of the InsP_3_R ion channel. The time needed to produce a manually-picked set precludes its adoption as a regular procedure for particle picking, and a method that reduced the temporal investment could prove valuable for researchers. In addition, scientists seeking particular idealized structures can consciously or subconsciously impart their own biases into manual picking, preferring certain angular views of the particle or omitting subsets of particles that do not exhibit anticipated features [5].

This work examines an increasingly popular method of data processing, crowdsourcing. A term coined in 2006, crowdsourcing opens a task normally assigned to a specific worker to a wider, more generalized userbase [6]. In recent years, crowdsourcing initiatives have come to rely on the ability of the internet to quickly disseminate data and recruit users to perform the necessary processing.

There are many approaches to crowdsourcing, including scientific games (e.g., Foldit, Eterna) and paid microtask services (e.g., Amazon Mechanical Turk, Crowdflower). Particularly intriguing is the emergence of ‘citizen science’ projects, which rely on community engagement and scientific intrigue to attract users to an otherwise menial task. Citizen science has proven extremely successful, with the project ‘Galaxy Zoo’ classifying over 1 million images from more than 100,000 users over nine months [7]. In this paper, we present and analyze the results from a citizen science project ‘Microscopy Masters’, which focused on crowdsourcing particle picking from single-particle cryo-EM micrographs. We examine the efficacy of crowdsourcing particle picking to lightly trained workers when compared to trained electron microscopists, and show that particle sets derived through crowdsourcing can yield robust and reliable 2D class averages and 3D reconstructions. The method presented here is shown as not only a viable time-saving option for datasets that confound automatic pickers, but also shows promise for future applications of crowdsourcing to cryo-EM data processing.

## 2. Results

### 2.1. Production of gold standard

A ‘gold standard’ or ‘ground truth’ for evaluating annotated subjects is crucial for beginning any classification study. In the case of algorithmic particle picking, evaluation is typically performed relative to a set of manually picked micrographs. Although manually picked datasets are available from previous studies examining particle picking, they generally contain a small number of images and are only annotated by a single individual [1]. In order to create a richer gold standard for evaluating our crowdsourcing protocol, micrographs were chosen from a single-particle cryo-EM dataset of the 26S proteasome lid complex [8]. Out of the 3,446 micrographs used in the published refinement, 190 were marked by at least two randomly-assigned cryo-EM experts, with a total of nine contributing participants.

In addition, intra-expert agreement was measured by requiring each expert to mark five randomly chosen images twice. The complete union of all marks by all experts totaled to 13,028 particles and was used as the ground truth for all following accuracy measurements in this paper.

Agreement between two annotations was calculated using the Jaccard index, defined as the proportion of the size of the intersection of the particles picked in two annotations to the size of the union. Intra-expert agreement was found to be surprisingly low and only slightly higher than inter-expert agreement, intra-agreement between all experts averaged 0.56 and inter-agreement averaged 0.45. This indicates that less than three quarters of particles picked by a single individual are picked again on reannotation by the same annotator on the same image. Among those annotators who completed all assigned micrographs, agreement was consistently higher among intra-expert comparisons (Table 1).

### 2.2. Initial Testing

A chief concern for crowdsourcing, especially citizen science, is building a reliable userbase, either through accessing pre-existing groups of users or attracting users through social media and community engagement.

**Table 1:** Intra-expert and inter-expert agreement for experts who created the gold dataset. Of the nine experts who participated, one did not complete all assigned micrographs and so was not included in this table.

| User | Inter | Intra |
| --- | --- | --- |
| Expert 1 | 0.45 | 0.64 |
| Expert 2 | 0.42 | 0.46 |
| Expert 3 | 0.54 | 0.65 |
| Expert 4 | 0.40 | 0.51 |
| Expert 5 | 0.44 | 0.64 |
| Expert 6 | 0.44 | 0.58 |
| Expert 7 | 0.45 | 0.61 |
| Expert 8 | 0.48 | 0.51 |

**Figure 1:**
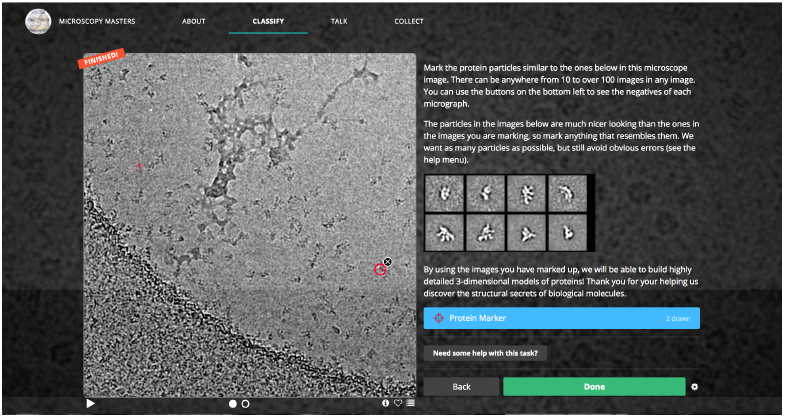
Screencap of picking interface hosted on Zooniverse

Since we desired to annotate a large, fully manually-picked particle set, we hosted our experiment on an established crowdsourcing platform, Panoptes, a Zooniverse-run initiative for citizen science projects.

Initial testing for the crowdsourcing system was performed by paid workers recruited through Amazon Mechanical Turk (AMT). Workers were recruited, trained, and paid through AMT, while Panoptes hosted the particle selection tasks and stored the results (Figure 1).

Testing produced 16,562 particles chosen by 42 unique workers using the same set of 190 images in the gold standard. Based off of feedback from AMT workers, additional instructions, shown in (Fig. S2), were added to the picking interface.

Importantly, this initial testing was used to determine the optimal number of people to assign to each image, as well as to establish a voting mechanism. To this end, at least 10 workers annotated each micrograph and accuracy statistics were derived for randomized subsets of those workers using various voting thresholds (Figure 2a). Limited returns after five annotators at union led to the choosing of this threshold as optimal; in all subsequent experiments, each image was shown to five annotators and the crowd output was defined as the union of users’ annotations.

### 2.3. Zooniverse

After testing in Amazon Mechanical Turk, the project, dubbed Microscopy Masters, was launched on Panoptes in March 2016. A total dataset of 209,696 particle picks was produced over a year from 3,446 micrographs, with 2,108 unique volunteers. The parameters established in our initial testing were utilized, with each image being classified by five different users and the ‘voting threshold’ set to one, meaning the total union of all classifications performed on an image were used to generate the final data set of picks. For individual users, we observed a marked decrease in fscore and recall in the Zooniverse set, as shown in Figure 2, which we attributed to differing incentives between paid testing on AMT and unpaid volunteers on Zooniverse. In particular, the number of particles selected in each image by Zooniverse volunteers is highly variable; a peak at zero in the distribution of particles picked per-user per-image resulted in a corresponding peak at zero for recall and fscore, as well as a peak at one for precision (Figure 2b). Association of low recall and low-cardinality annotations implied a body of “low-effort” annotations, where a user did not fully complete the image before submission. Aggregation of the five user annotations per image mitigated the low individual accuracy, yielding an average aggregate fscore comparable to that of the AMT-annotated data, as shown in (Figure 2d).

**Figure 2:**
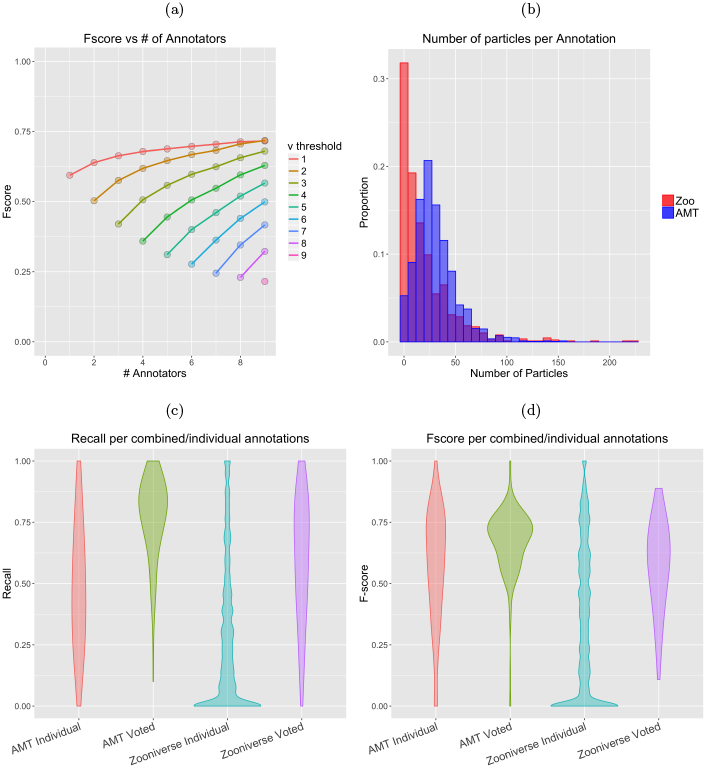
(a) Number of annotators vs performance, grouped by voting threshold. The top line represents union, while the bottom rung represents consensus. (b) Histogram showing distribuion of number of particles picked per person per image for both AMT and Zooniverse. Note peak at zero for Zooniverse, comprising ‘low-effort’ annotations. Recall (c) and F-score (d) of individual classifications from Zooniverse and AMT workers compared to gold standard. High numbers of low-quality, low-effort annotations from Zooniverse workers result in peaks at 0 for recall and fscore.

Average agreement between the voted crowd annotations and individual expert annotations was found to be slightly less than inter-expert agreement, with the mean for inter-expert agreement at 0.45 and mean between the crowd and experts at 0.40 (Figure 3).

**Figure 3:**
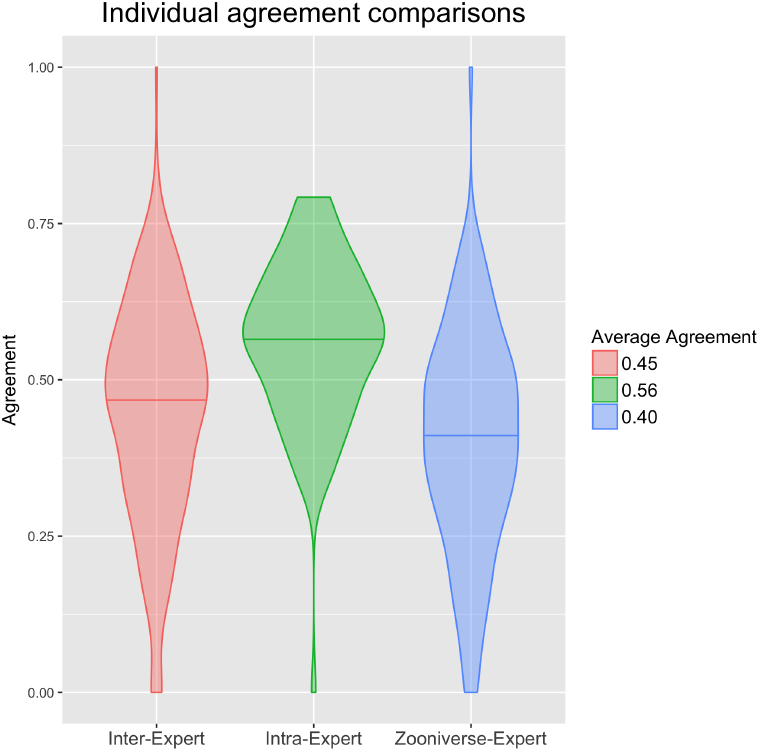
Violin plots of agreement for inter-expert, intra-expert, and crowd-expert comparisons. Standard deviations (σ) and number of comparisons (*n*) for each distribution are as follows: Inter-Expert: *σ* = 0.18, *n* = 227; Intra-Expert: *σ* = 0.13, *n* = 90; Zooniverse-Expert: *σ* = 0.18, *n* = 382.

### 2.4. Reconstruction

Refinement of the proteasome lid complex structure was performed using the crowdsourced dataset collected through Zooniverse, as well as the dataset used in Dambacher et al., which was picked using a template-based method [8]. Since the resolution of a refined single-particle reconstruction generally correlates with the number of particles in the dataset, a reconstruction was also generated using a random subset of the template-picked particles with the same cardinality as the crowdsourced data, called the normalized template set [9]. Particle stacks were extracted from the micrographs, and 2D reference-free classification was performed. “Junk classes containing false particle picks or damaged/aggregated particles were manually selected and removed, following the same selection criteria used previously [8]. A subset of homogeneous high resolution particles were identified through 3D classifications which were refined to yield the final reconstruction (Figure 4). The number of particles remaining after each filtering step, as well as the final resolution estimates, indicate the template sets produced reconstructions with more internal structural consistency (Table 2). FSC curves for each reconstruction, used for resolution estimation, indicate no irregularities in any of the particle sets (Figure 5). Resolution was found to be lower for the refinement produced from the automatically picked datasets, with the Zooniverse-produced refinements resolution at 4.148 ˚A and the template-picked refinements resolution at 3.829 ˚A as measured by FSC curves at FSC= 0.143. The refinement produced from the normalized template set was 4.036 ˚A, predictably higher than the template refinement and slightly lower than the crowdsourced refinement. As a further measure of resolution consistency, ten atomic models were independently built into the reconstructions [10]. The root-mean-square deviation (RMSD) of each residues Cα position in the ten reconstructions was used as a measure of the local quality of the refinements, with a higher RMSD indicating a less convergent refinement (Figure 6). While the distribution of RMSD values from the crowd set were significantly higher than the template-picked structures (p < 0.01 as assessed by a two-sample Kolmogorov-Simrnov test), the overall magnitude of the difference was relatively small. The normalized and full template reconstructions had 67.3% and 69.6% of residues with high convergence, respectively, while the crowd-produced reconstruction had 63.7% of residues with high convergence.

**Figure 4:**
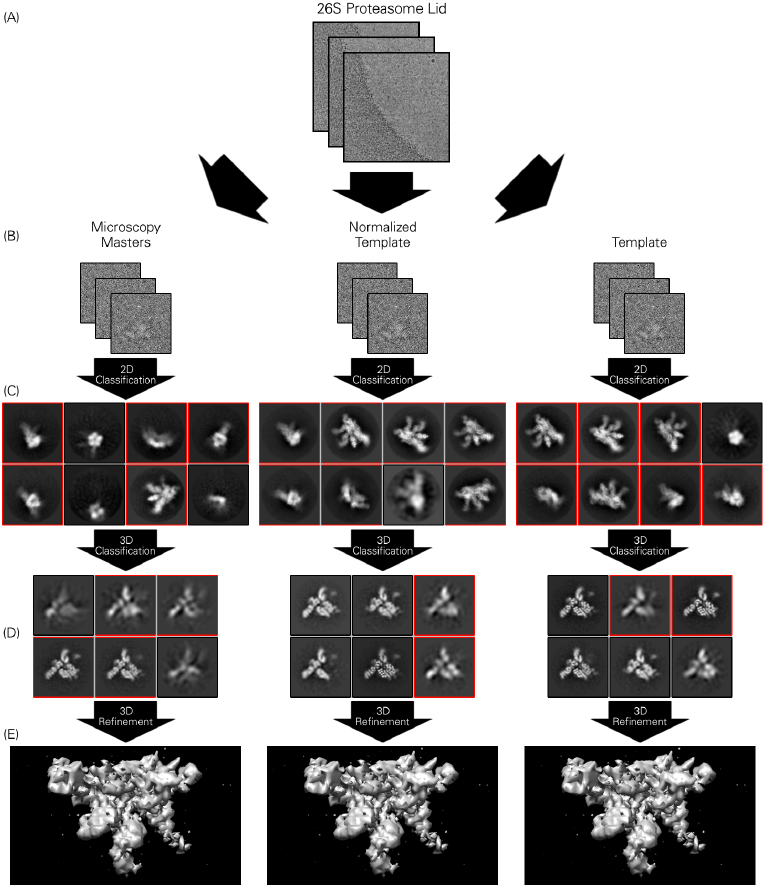
Reconstruction using crowdsourced, template-picked, and ‘normalized’ template-picked particle sets. (A) Micrographs from 26S are run through template and crowdsourcing particle picking procedures. (B) Particles are then extracted and run through 2D template-free class averaging. (C) Top 8 2D classes from each particle set. Classes chosen to continue through processing marked in red. (D) All 3D classes from 3D classification of particles chosen in step (C), with similar marking. (E) Final refinements for each particle set.

**Figure 5:**
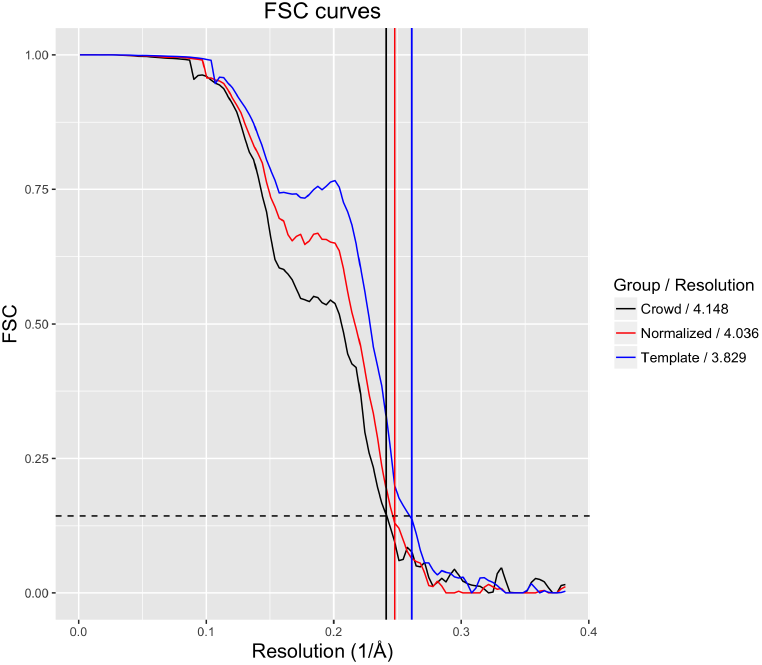
FSC curves for all reconstructions. Horizontal dash line at *y* = .143 represents value for resolution estimation. Vertical lines’ colors correspond to FSC curves and the x-intercepts represent each curve’s resolution.

**Figure 6:**
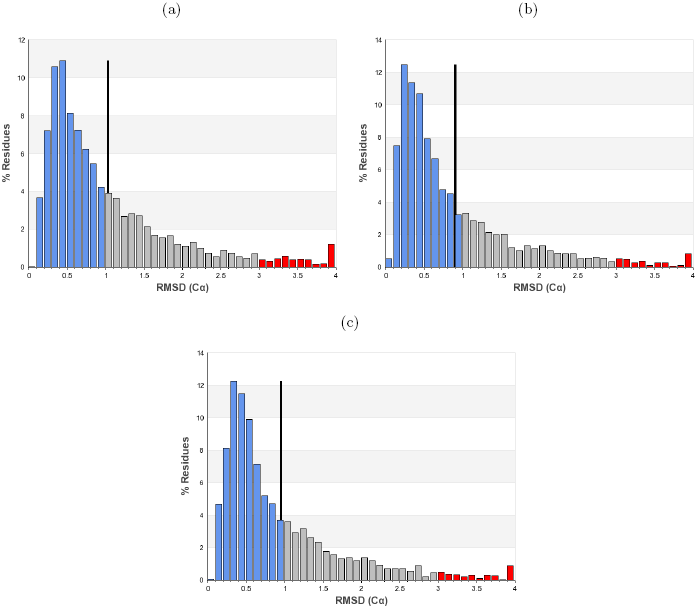
RMSD histograms of Cα positions in the tenfold ensemble analysis. Blue, gray and red bars denote residues of low ( < 1), medium(> 1,< 3, and high (> 3) convergence, respectively. The black line marks the mean of the distribution. (a) Ensemble based off refinement from Zooniverse volunteers. (b) Ensemble from automated template-based method data. (c) Ensemble from normalized automated template-based method data.

## 3. Discussion

Our study has demonstrated the utility of crowdsourcing particle picking for single-particle cryo-EM to users with little to no experience in cryo-EM. Volunteer particle pickers created a sizeable, usable dataset through a popular citizen science site that produced a 3D reconstruction of comparable quality to the reconstruction generated from template-picked particles. The particles chosen by citizen scientists also compared favorably to those produced in paid scenarios, suggesting the use of a citizen science framework as a low-cost, low-effort alternative to producing high-quality datasets without the potential biases associated with template picking or manual selection by a single person.

**Table 2:**
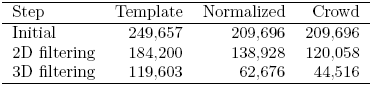
Number of particles left after each filtering step during single-particle reconstruction.

| Step | Template | Normalized | Crowd |
| --- | --- | --- | --- |
| Initial | 249,657 | 209,696 | 209,696 |
| 2D filtering | 184,200 | 138,928 | 120,058 |
| 3D filtering | 119,603 | 62,676 | 44,516 |

Many automatic picking algorithms still rely on some amount of manual picking, since popular template-driven algorithms generally use 2D class averages generated from manually-picked particles as templates. The demonstrated lack of consistency of scientists trained in single-particle cryo-EM when manually picking calls into question the acceptance of manually-picked data as the ground truth when assessing particle-picking protocols. More generally, the noisy nature of micrographs can make it difficult for the human eye to reliably distinguish the multitude of patterns needed to create a robust single-particle dataset, particularly for smaller-sized complexes.

Key to the acceptance of manual selection as a blue-chip particle-picking method is the superior quality of resultant reconstructions. In our trials, reconstructions produced from the full and normalized template sets, as well as the manually picked set, showed no marked differences in resolution. Far more particles were filtered from the manually picked set through classification, likely a result of the template consistently picking particles with similar features, which results in better clustering of 2D classes. However, since manual picking does not rely on templates, 2D views of protein particles not present in the templates could be recovered from micrographs.

Manual picking through crowdsourcing approaches can produce a dataset of comparable quality to both manual picking by trained experts and computational particle picking methods, but the question remains as to whether manually picked data produces significantly better reconstructions.

Volunteer particle pickers produced a large body of “low-effort” annotations when they were asked to work for no pay. The deleterious effects of these annotations are successfully mitigated by using the union of particles from five separate annotations for each image. Since the low-accuracy annotations generally are those with little to no marks, the increased accuracy from combining all annotations can be inferred to come from users who contribute particles that generally agree with experts’ picks. However, relying upon a fraction of annotations is not ideal, and future attempts to crowdsource particle picking should focus on creating incentive structures that encourage a higher proportion of high-quality annotations. A further limitation to crowdsourcing is the time needed to produce a manually picked dataset, with our project running over the course of a year before completion. Producing a fully crowdsourced particle set might be most effective for proteins where the particles are particularly difficult to identify with template pickers or other automatic methods.

As mentioned previously, automated template particle pickers generally require some initial manual selection of particles from micrographs in order to to ‘seed’ the algorithm with 2D templates of the desired protein particle. In order to explore the potential for crowdsourcing in accomplishing the preliminary manual task of generating such seed datasets of particle picks, three datasets were presented to untrained workers with the intent of creating templates suitable for automated particle picking programs. A small number of micrographs from datasets of the HIV-1 envelope trimer, and TRPV2 ion channel were presented to Zooniverse and AMT users [11, 12].

It took two days for the users to complete the desired one hundred HIV-1 trimer annotations, and eight days for the desired one hundred classifications, the discrepancy likely caused by stagnation in the userbase of Microscopy Masters. On AMT, users completed the TRPV2 ion channel tasks in under an hour, while the HIV trimer tasks, which offered much less pay, took around two weeks. The same classification aggregation criteria used in the full reconstruction were applied to to the sprint datasets, yielding sets of 1863 and 3298 particles for the HIV-1 trimer and TRPV2 datasets, respectively. Reference-free 2D classification of the manually selected datasets yielded averages that were suitable for use as templates (Fig. S4). Assuming an engaged group of users can be accessed, either through a paid system or by cultivating a dedicated community, the relatively short turnaround time of creating templates via this method demonstrates its usefulness for cryo-EM researchers.

Another possible application for crowdsourcing can be found in cellular tomography, which requires manual curation in order to identify and link sub-cellular structures in multiple frames. Even newer machine-learning algorithms for automated segmentation require that a user initially identify features manually within tomograms [13].

The results outlined in this manuscript showcase crowdsourcing as a useful, new option for microscopists whose data resist selection by automated method, but although the untrained workers demonstrated an ability to produce particle sets of acceptable quality, there is work to be done in creating viable incentive structures and encouraging user engagement. Despite the difficulties in producing a fully crowdsourced dataset, the body of work presented here is an encouraging first step in the novel application of crowdsourcing processing for cryo-EM data.

## 4. Methods and Materials

### 4.1. Gold Standard Production

The gold standard was generated in a single, pizza-fueled afternoon with resident electron microscopy researchers at The Scripps Research Institute. Each researcher was randomly assigned forty-five images out of a 190 micrograph subset from the 26S Proteasome lid complex dataset from Dambacher et al. [8]. Additionally, each expert marked five images twice in order to study consistency of experts’ particle picking, making a total of fifty annotations for each expert. The union of these repeated images’ picks were used for the gold standard of a particular expert on the respective image, and the union of all experts’ picks was used as the gold standard for our entire study. The platform used by the experts was identical to that used by the AMT workers, although some features such as the pop-up tutorial and additional screen for marking unusable images implemented in the Microscopy Masters project were absent for the experts.

### 4.2. Project hosting on Zooniverse

Initiation of our crowdsourcing project coincided with the launch of the Panoptes system on a popular crowdsourcing consortium called Zooniverse, creators of the successful Galaxy Zoo. Panoptes offered a customizable interface for allowing the Zooniverse userbase to actively work on projects outside of the usual Zooniverse scope. Since the Zooniverse userbase numbers in the millions, its use offered access to a much larger group of users than could have been achieved on an independent platform.

### 4.3. Initial user testing

In order to create a usable, easily understandable interface, the Panoptes workflow was initially tested on Amazon Mechanical Turk (AMT). Workers recruited via AMT were sent to single images on Panoptes and given a code unique to each image to verify that they completed the task. Before being allowed to work, completion of a short tutorial and mutiple-choice quiz was required, with a minimum allowable score of 5/7 questions correct. Initially, instructions beyond initial quizzing and explanation were minimal, but this led to complaints from the userbase that it was too difficult to complete the task without any easily accessible instructions. Workers were paid variable amounts from ten cents to 25 cents per image, they generally averaged 20 seconds per image. Predictably, lower payments resulted in a lower rate of participation; 10 cent payments took one week to complete at five annotations per image over 200 images, while 25 cent payments with the same amount of required work took 3 days. Text for the tutorials and help text were tweaked using feedback from the AMT workers before a full launch on Panoptes.

### 4.4. Microscopy Masters

Initial testing was followed by a beta release of Microscopy Masters to select members of the Zooniverse community. Following feedback from the beta test, an additional screen allowing users to mark images as poor quality was placed before the picking interface. Some features were added to the interface during the middle of production; two versions were implemented over the course of the project. When compared to gold standard, the differing versions showed no marked dissimilarity in terms of fscore.

### 4.5. Voting Protocol

Although the ‘voting threshold’ for points was set to one, meaning that a point only needed to be selected once for it to be included in the final set, an algorithm needed to be designed to combine picks from separate annotators. Two points in two separate annotations of the same image were combined into a vote if:

1. They were within 20 pixels of one another
2. They were mutual nearest neighbors, i.e., each point was the other’s nearest neighbor in the second annotation

If two points were combined, their averaged coordinates were used as a new particle.

### 4.6. 26S Proteasome lid complex dataset

A single-particle cyro-EM dataset for the ovine 26S proteasome lid complex was chosen as the primary test case for crowdsourced particle picking. The globular nature of the lid complex can confound often-used template pickers, making it an ideal choice to test the limitations of automatic picking procedures. Dambacher et al. imaged the dataset on a Krios Titan cryo-EM microscope, technical details are detailed in their text [8].

### 4.7. Reconstruction Pipeline

Cryo-EM processing was performed in Relion v1.4 [14]. Processing followed the methods previously described [8] in order to produce comparable reconstructions. CTF parameters were estimated using CTFFIND3 [15]. Particles were extracted from respective particle coordinate files with a box size of 256 pixels. Choosing of class averages was performed through random selection by Gabriel Lander; classes from separate reconstructions were anonymously mixed treated as a single dataset. Particles from chosen classes were passed on to the subsequent step. Final refinement was performed with the same initial model used in Dambacher et al. using default parameters in RELION.

### 4.8. Initial Template Construction

Images for the initial template construction were initially processed in a manner similar to that of the large 26S proteasome lid project. Five classifications were requested for the HIV trimer set on both platforms as an initial test, this was increased to fifteen classifications for TRPV2 on zooniverse in order to attract more users for annotation. In an attempt to make the task more visually engaging, micrographs were colored with various contrasting colors as well as offering the traditional black and white. No marked decrease in quality was associated with the color changes. Upon release of the new datasets, Microscopy Masters users were notified of additional datasets via an emailed newsletter.

## 4.9. Acknowledgements

We thank the members of the Hazen cryo-EM facility for their help in generating the gold standard dataset. We also thank J.C. Ducom at the High Performance Computing group for computational support. G.C.L. is supported as a Searle Scholar, a Pew Scholar, and by the National Institutes of Health (NIH) DP2EB020402. This work was also supported by the National Institutes of General Medical Sciences (U54GM114833 to A.I.S.).

---

**Figure S1:**
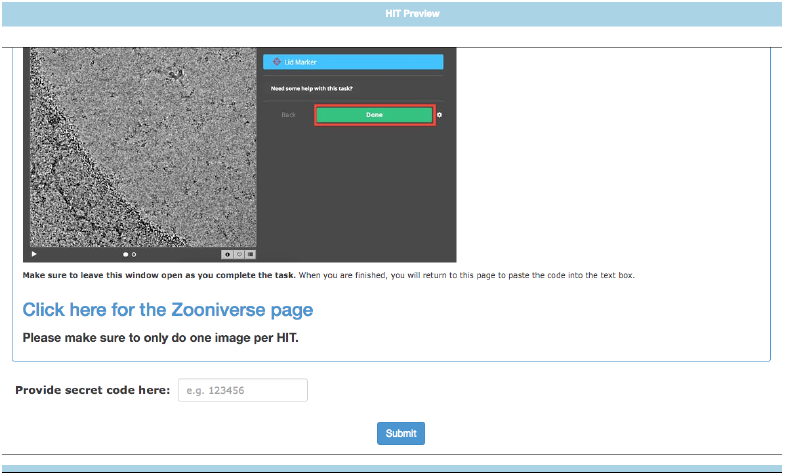
Screencap of AMT HIT page

**Figure S2:**
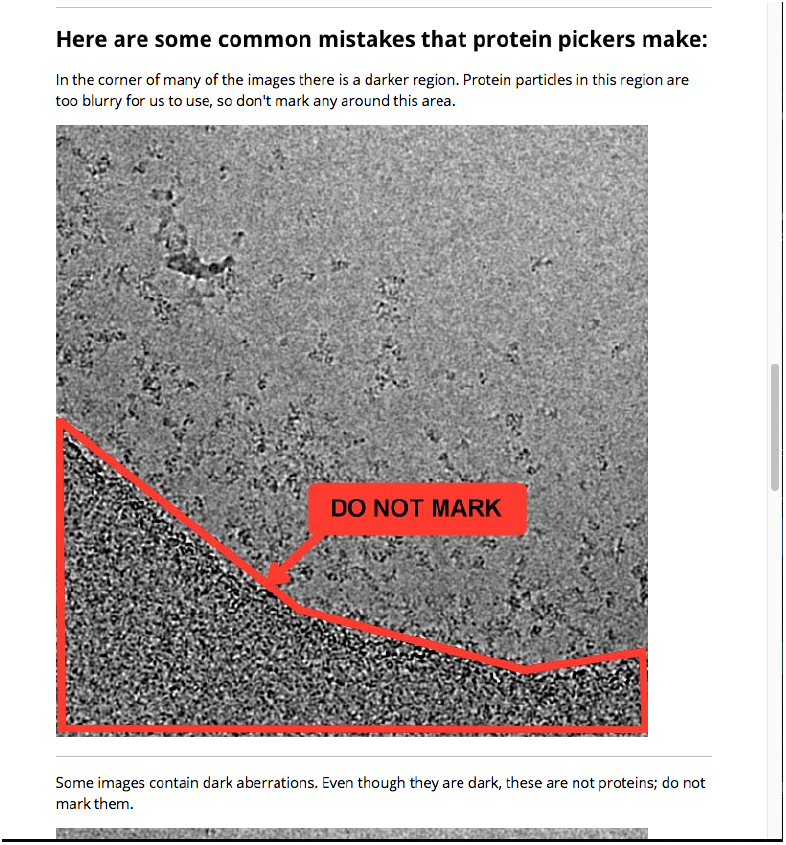
Example of instructions given to Zooniverse and AMT workers on incorrect particle picking

**Figure S3:**
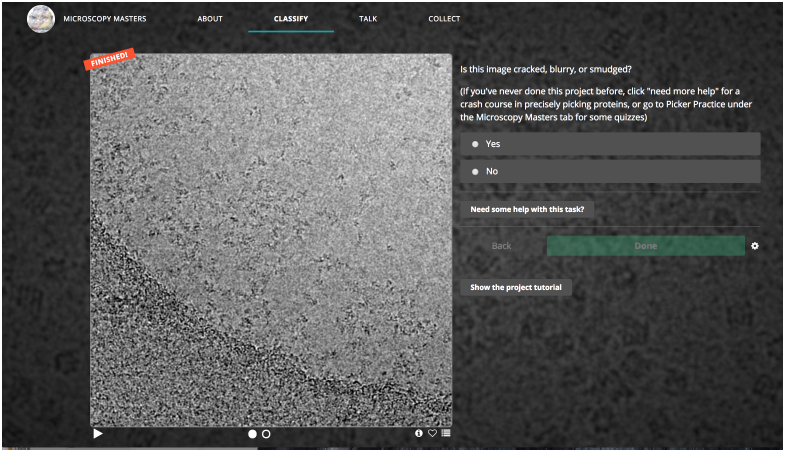
Screencap of Initial Zooniverse page

**Figure S4:**
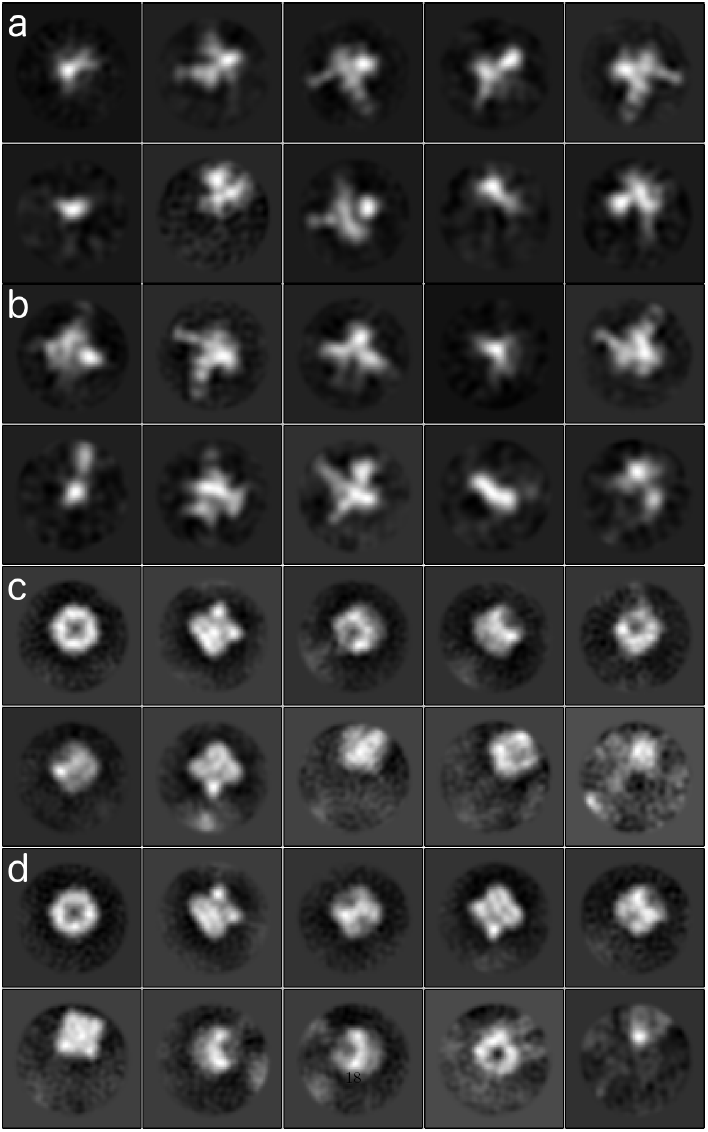
Top 10 2D class averages from each of the initial template construction experiments. (a) & (b) show results from AMT and Zooniverse, respectively, of the HIV envelope trimer. (c) & (d) show results from AMT and Zooniverse, respectively, of the TRPV2 ion channel.

